# Connexin50 hemichannels are opened by CO_2_: implications for lens physiology

**DOI:** 10.1101/2025.01.23.634273

**Authors:** Alexandra Lovatt, Frederick Bibra, Ashvini Wijayapala, Macy Mui, Jack Butler, Nicholas Dale

**Author notes:** Contributed equally (Alexandra Lovatt, Frederick Bibra).

## Abstract

Connexin50 (Cx50) is expressed in lens fibre cells. As mutations of Cx50 cause cataracts its physiological role in the lens must be important. We have used recent cryoEM structures of Cx50 and the predictive power of Alphafold3 to identify the presence of a carbamylation motif, originally described in Cx26, that suggests that Cx50 might be CO_2_ sensitive. By expressing the naturally truncated version of Cx50 in HeLa cells and utilising coexpression of genetically encoded sensors iGluSnFr or eLACCO1.1, we have demonstrated the CO_2_-dependent opening of Cx50 hemichannels, and their permeability to lactate and glutamate. By mutating the two key residues of the carbamylation motif, K105 and K140, we have shown that the motif is required for CO_2_ sensitivity. Mutations of the residue V44 cause cataracts and these mutations abolish the CO_2_ sensitivity of Cx50. Using Fluo-4 Ca^2+^ imaging with lens slices we have demonstrated CO_2_-dependent Ca^2+^ influxes into fibre cells that are blocked by La^3+^ and exhibit the same CO_2_ dose dependence as Cx50 hemichannels. Lens fibre cells respond to glutamate via NMDA receptors and our data shows that the Ca^2+^ influx to raised PCO_2_ partially depends on NMDA receptor activation. We hypothesize that CO_2_-dependent gating of Cx50, subsequent release of glutamate resulting in the downstream activation of glutamate receptors, and the consequent alterations in transmembrane Na^+^ fluxes, provide homeostatic control of the microcirculation system that is critical for lens health.

## Introduction

Connexin50 (Cx50) is an alpha connexin that shares considerable sequence homology and structural similarity to Cx46 (Myers *et al*., 2018; Flores *et al*., 2020). Both Cx50 and Cx46 are part of the alpha connexin clade which also includes Cx43 (Cruciani & Mikalsen, 2006). These three connexins are all expressed in the mammalian lens (Mathias *et al*., 2010; Berthoud & Ngezahayo, 2017). While Cx43 is found only in the lens epithelium, Cx46 and Cx50 form gap junctions between the lens epithelial and lens fibre cells, and between lens fibre cells. Cx50 is also present in lens fibre cells in the form of hemichannels (Shi *et al*., 2018; Wang *et al*., 2025) -unopposed hexameric channels that open into the extracellular space.

To maintain transparency, the lens does not have a blood supply (Mathias *et al*., 1997). Essential nutrients and metabolic waste products are supplied/removed via a microcirculation system (Mathias *et al*., 1997; Mathias *et al*., 2007; Delamere & Tamiya, 2009). This is powered by Na^+^ extrusion from equatorial epithelial cells. The movement of Na^+^ ions through the gap junction coupled network of lens cells drags along water and other solutes. Water and these solutes re-enter the lens at the poles through the expanded extracellular space provided by the lens sutures.

An open question remains the pathway by which Na^+^ ions enter the lens fibre cells (Mathias *et al*., 1997) to generate extrusion via the Na^+^/K^+^ pumps in the epithelial cells and thus the microcirculation. Cx50 has been discounted as the source, because the hemichannels are deemed to be closed and instead an unspecified leak channel has been proposed (Mathias *et al*., 2007). Hemichannels of Cx46 have been proposed to contribute to this leak current (Ebihara *et al*., 2014). In this paper we show that Cx50 hemichannels are directly sensitive to gaseous CO_2_ which acts via a binding mechanism and motif that we have previously shown in Cx26 (Meigh *et al*., 2013). In Cx26, CO_2_ forms a covalent carbamate bond with Lys125 (Nijjar *et al*., 2025). The subsequent carbamate is negatively charged and can interact with Arg104 of the neighbouring subunit. The carbamate bridges are thought to trap the Cx26 hemichannel in the open configuration (Brotherton *et al*., 2022; Brotherton *et al*., 2024). Cx26 is not unique in its CO_2_ sensitivity, hemichannels of Cx30 (Huckstepp *et al*., 2010a), Cx32 (Huckstepp *et al*., 2010a; Dospinescu *et al*., 2019; Butler & Dale, 2023) and most recently Cx43 (Dospinescu *et al*., 2025) are also opened by CO_2_ acting via equivalent motifs.

The properties of Cx50 hemichannels show that they will be partially open at levels of PCO_2_ that are typical of aqueous humour (Krupin *et al*., 1980). This suggests that Cx50 hemichannels have the correct properties to be the source of the Na^+^ leak current that underlies the microcirculation. By using Ca^2+^ imaging with acute lens slices and varying PCO_2_ we have provided some evidence that supports this hypothesis.

## Materials and Methods

### Connexin mutagenesis

A cDNA sequence for the human gene for Cx50 according to accession P48165 truncated to 277 amino acids was synthesised by IDT. This truncation of the human gene is equivalent to the natural age-related truncation at His284 documented for the bovine gene (Wang & Schey, 2009). The cDNA sequence was subsequently subcloned into pCAG-GS-mCherry vector prior to transfection. Point mutations were introduced via Gibson assembly methods. Overlapping primer fragments (IDT) both containing the desired mutation were amplified via PCR. Successful mutagenesis was confirmed using Sanger sequencing (GATC Biotech). All Cx50 constructs were inserted upstream of an mCherry tag, linked via a 12 AA linker (GVPRARDPPVAT).

### Cell culture and transfection

Parental HeLa DH cells (ECACC 96112022, RRID:CVCL_2483) were cultured with low-glucose DMEM (Merck Life Sciences UK Ltd, CAT# D6046) supplemented with 10% foetal bovine serum (Labtech.com, CAT# FCS-SA) and 5% penicillin/streptomycin. Cells were seeded onto coverslips at a density of 4×10^4^ cells per well. Cells were transiently transfected to co-express a Cx50 variant and one of the following genetically encoded fluorescent sensors:

pCMV(MinDis).iGluSnFR was a gift from Loren Looger (Addgene plasmid # 41732; http://n2t.net/addgene:41732; RRID:Addgene_41732) (Marvin *et al*., 2013).

pAEMXT-eLACCO1.1 was a gift from Robert Campbell (Addgene plasmid # 167946; http://n2t.net/addgene:167946; RRID:Addgene_167946) (Nasu *et al*., 2021). To improve expression of eLACCO1.1, this construct was subcloned into the iGluSnFR expression vector backbone. Sequences were verified with Sanger sequencing (GATC).

To transfect cells, a mixture of 1μg of DNA from pCAG-Cx-mCherry construct and 1μg sensor DNA with 3μg PEI was added to cells for 4-8h. Cells were imaged 48h after transfection.

### Recording solutions used

20 mmHg PCO_2_: 140 mM NaCl, 10 mM NaHCO_3_, 1.25 mM NaH_2_PO_4_, 3mM KCl, 1 mM MgSO_4_.

35 mmHg PCO_2_: 124 mM NaCl, 26 mM NaHCO_3_, 1.25 mM NaH_2_PO_4_, 3mM KCl, 1 mM MgSO_4_.

55 mmHg PCO_2_: 100 mM NaCl, 50 mM NaHCO_3_, 1.25 mM NaH_2_PO_4_, 3 mM KCl, 1 mM MgSO_4_.

70 mmHg PCO_2_: 70 mM NaCl, 80 mM NaHCO_3_, 1.25 mM NaH_2_PO_4_, 3 mM KCl, 1 mM MgSO_4_.

High K^+^ (20 mmHg PCO_2_): 93 mM NaCl, 10 mM NaHCO_3_, 1.25 mM NaH_2_PO_4_, 50mM KCl, 1 mM MgSO_4_.

10 mM D-glucose and 2 mM CaCl_2_ was added to all solutions just before use. Solutions were and saturated with: 98%O_2_/2% CO_2_ (20 mmHg); 95% O_2_/5% CO_2_ (carbogen) (35 mmHg), or carbogen plus addtional CO_2_ (55 mmHg and 70 mmHg). The amounts of CO_2_ were adjusted so that all solutions had a pH of ∼7.4.

### Live cell fluorescence imaging and analysis

48 hours after transfection, cells were perfused with control aCSF (20 mmHg) until a stable baseline was reached, before switching to either hypercapnic or high K^+^ aCSF. Once a stable baseline was reached after solution change, cells were returned to perfusion with control aCSF (20 mmHg). Recordings were calibrated by application of 3 μM of the corresponding analyte.

All cells were imaged by epifluorescence (Scientifica Slice Scope, Cairn Research OptoLED illumination, 60x water Olympus immersion objective, NA 1.0, Hamamatsu ImagEM EM-CCD camera, Metafluor software). cpGFP in the sensors were excited by a 470 nm LED, with emission captured between 504-543 nm. Connexin constructs have a C-terminal mCherry tag, which was excited by a 535 nm LED and emission captured between 570-640 nm. Only cells expressing both cpGFP and mCherry were selected for recording, with cpGFP images acquired every 4 seconds. For each condition, at least 3 independent transfections were performed with at least 2 coverslips per transfection.

Analysis of all experiments was carried out in ImageJ. Images were opened as a stack and stabilised for movement (Li, 2008). ROIs were drawn around cells co-expressing both sensor and connexin. Median pixel intensity was plotted as normalised fluorescence change (ΔF/F_0_) over time to give traces of fluorescence change. The amount of analyte release was quantified as concentration by normalising to the ΔF/F_0_ caused by application of 3 μM of analyte, which was within the linear portion of the dose response curve for each sensor.

### Membrane localisation of Cx50 and mutants

HeLa cells adhered to coverslips were transfected with mCherry-tagged wild type and mutant versions of Cx50. After 48h, the cells were washed 3 times with PBS, fixed with 4% paraformaldehyde in PBS for 30 mins. They were then washed 3 times with PBS and incubated with serum free DMEM containing 2.5 μM DiO (3,3′-Dioctadecyloxacarbocyanine perchlorate, Sigma-Aldrich Cat#D4292) for 15 minutes. After three further washes in PBS, coverslips were mounted on glass microscope slides in a Fluorshield™ with DAPI mounting medium (Sigma-Aldrich, Cat# F6057). They were then imaged on a Zeiss 980 LSM confocal microscope using the 488 and 561nm excitation wavelengths for DiO and mCherry respectively.

Colocalization analysis between the mCherry tag of the Cx50 variants and the DiO membrane stain was performed with Fiji and the JaCoP plugin (Bolte & Cordelieres, 2006). ROIs were drawn around mCherry-positive cells and the image surrounding the ROI was removed. The Manders’ co-efficient (Manders *et al*., 1993) was used as a quantitative assessment of colocalisation, and hence the membrane localisation of the Cx50 variants. The images were thresholded to include DiO membrane staining but exclude any diffuse background fluorescence.

### Dye loading of whole lens

Live, whole lens was harvested and immediately perfused with control (20 mmHg) aCSF. Upon experimentation, samples were transferred to either control, hypercapnic or hypercapnic aCSF with 200 µM LaCl_3_ (Sigma-Aldrich, Cat #10025-84-0), each containing 50 µM fluorescein isothiocyanate (FITC) (Sigma-Aldrich Cat #46950) for 10 mins. Samples were then perfused with FITC-containing control aCSF for 20 mins, preventing dye leakage.

Lenses were then washed serially in control aCSF and fixed in 4% paraformaldehyde (PFA) for 2 h at room temperature. Fixed tissue was washed at room temperature in PBS. The tissue was then embedded 4% PBS-based agarose gel and mounted in an ice-laden sectioning chamber with. Slices were cut either coronally or transversely to 105 µm on a vibratome (Leica VT1200), and arranged serially in 24-well plates, stored in PBS. Free-floating sections were then treated with 20 µM DiI (Sigma-Aldrich, Cat #468495), a lipophilic membrane stain, for 30 mins at room temperature. These samples were washed and mounted upon polysine-coated slides (Polysine, VWR) and dehydrated at room temperature for 10 mins. Coverslips were applied using FluorshieldTM with DAPI mounting medium (Sigma-Aldrich, Cat #F6057). Specimens were imaged with a Zeiss-980 confocal LSM under a 63X oil immersion lens, using 488 the 561 lasers, using the same settings across all slices imaged to allow direct comparison of fluoresence intensity.

Images were again analysed in FIJI, blind to perfusion condition. ROI’s were drawn around individual cells, measuring mean pixel intensity and thus dye loading extent. Measurements were calibrated by subtracting the mean pixel intensity of a representative background ROI within the same image. A minimum of 5 lenses were included per condition, with a mean value of the fluorescence from at least 10 cells used as the data point.

### Preparation of lens slices

Live whole lenses were incubated for 18 mins in decapsulation solution first described by Dewey et al (1995): 149.2 mM KMeSO_3_, 5 mM HEPES, 2 mM EGTA, 2 mM EDTA and 5 mM glucose and included 0.10 mM bumetanide (diluted from a 10 mM stock solution in absolute ethanol). This treatment allowed separation of the capsule with minimal teasing.

Lenses were individually embedded in 10% ultra-low gelling temperature agarose (Sigma-Aldrich, Cat # A5030) made in 20 mmHg Ca^2+^-free aCSF, placed in an ice-laden cutting chamber, submerged in 20 mmHg Ca^2+^-free aCSF including 1 mM kynurenic acid, and sectioned either coronally or transversely, using a vibratome to create a single central slice 300 µm thick.

### Ca^2+^ imaging of lens slices

Fluo4-AM (Invitrogen, Cat #F14201) dissolved in Pluronic™ F-127 (Invitrogen, Cat #P3000MP) with the aid of sonication and vortexing was diluted into 20 mmHg aCSF to a final concentration of 2.5 μM. Lens slices were incubated for 25 mins in a custom humidified perfusion microchamber, with superperfusion of 98% O_2_/2 % CO_2_ to maintain pH and PO_2_, before a subsequent 25 min wash with 20 mmHg aCSF.

Loaded lens slices were placed in a perfusion chamber and subjected to control aCSF until a stable baseline was maintained. Changes in Fluo-4 fluorescence intensity from baseline, defined as ΔF/F_0_, were measured via epifluorescence (Scientifica Slice Scope, Cairn Research OptoLED illumination, 60x water Olympus immersion objective, NA 1.0, Hamamatsu ImagEM EM-SSC camera, Optofluor software). Fluo-4 was excited using 470 nm LED, with fluorescence emission recorded between 507 and 543 nm. Analysis of Ca^2+^ signals was performed in FIJI. ROIs were drawn around individual cells or regions, with tissue movement mitigated using the image stabiliser plugin. A recording from a unique lens was considered an individual replicate.

### Statistical analysis

For the iGluSnFr and eLACCO1.1 recordings, each cell was considered as an independent replicate. Pairwise comparions of the iGluSnFr and eLACCO1.1 recordings were made with the Mann Whitney *U*-test. Dye loading data, where multiple comparisons were made, was analysed with the Kruskal Wallis ANOVA and post hoc Mann Whitney *U*-tests. The Friedman 2-way ANOVA was used to analsye the Fluo-4 traces as multiple manipulations were performed on the same lens slice thus the lens slice was one factor and the pharmacological manipulation was the second factor. In this case post hoc testing was performed via the Wilcoxon matched pairs signed rank test. All quantitative data is presented as box and whisker plots, where the line represents the median, the box is the interquartile range, and the whiskers are the range, with all individual data points included. All calculations were performed on GraphPad Prism.

## Results

### Cx50 hemichannels possess the carbamylation motif and are CO_2_ sensitive

Alignment of the Cx50 sequence with that of Cx26 and Cx43, both of which are CO_2_ sensitive (Huckstepp *et al*., 2010a; Meigh *et al*., 2013; Dospinsecu *et al*., 2025) revealed the presence of the carbamylation motif _140_KFRLEGT_146_ in Cx50. This sequence is analogous to the motifs of _144_KVKMRGG_150_ in Cx43 and _125_KVRIEGS_131_ in Cx26. The carbamylation motif is within the cytoplasmic loop, which has proven to be notoriously diticult to resolve in cryoEM structures for all connexins (Maeda *et al*., 2009; Myers *et al*., 2018; Flores *et al*., 2020; Lee *et al*., 2020; Brotherton *et al*., 2022; Qi *et al*., 2023; Brotherton *et al*., 2024). Experimental Cx50 structures (e.g. 7JJP) show that K105 is in a similar orientation as K105 of Cx43 and R104 of Cx26. These latter residues form a salt bridge with the carbamylated Lys residue from the neighbouring subunit. Understanding the location of K140 is more difficult. The alphafold3 prediction for the cytoplasmic loop and the position of K140 is of low confidence and experimental structures do not resolve the entire cytoplasmic loop. However, some structures (e.g. 7JM9), while not resolving K140, do place R142 close to its location in the alphafold3 prediction suggesting that the predicted positioning of K140 is likely to be approximately correct. Therefore, there is a reasonable probability that K140 is sufficiently close to K105 to form an inter-subunit salt bridge following carbamylation of K140.

As Cx50 appears to have the carbamylation motif, we tested its CO_2_ sensitivity by expressing Cx50 in HeLa DH cells. To assay hemichannel opening we measured the possible efflux of either glutamate or lactate via Cx50 by co-expressing the genetically encoded sensors iGluSNFR or eLACCO1.1. We used these sensors as opposed to the more obvious use of GRAB^ATP^ to assay ATP release (Butler & Dale, 2023), as we found in exploratory experiments that Cx50 hemichannels are not permeable to ATP (Lovatt et al 2025).

Parental HeLa cells that expressed only iGluSnFr or eLACCO1.1 did not exhibit fluorescence changes in response to hypercapnic and depolarising stimuli (Fig 1). To test the possible CO_2_-sensitivity of Cx50 hemichannels, we next co-expressed this connexin with each of the genetically encoded sensors. Starting at a PCO_2_ of 20 mmHg, we successively switched to 35, 55 and 70 mmHg PCO_2_ solutions. We observed CO_2_-dependent fluorescence changes for both analytes (Fig 2). We therefore conclude that Cx50 hemichannels are CO_2_ sensitive and will permit CO_2_-dependent efflux of glutamate and lactate. Interestingly, significant release of glutamate and lactate was observed at a PCO_2_ of 35 mmHg. Maximal glutamate and lactate release was observed at a PCO_2_ of 55 mmHg and some inhibition of release, relative to this maximum, was observed at a PCO_2_ of 70 mmHg (Fig 2C,D).

**Figure 1.**
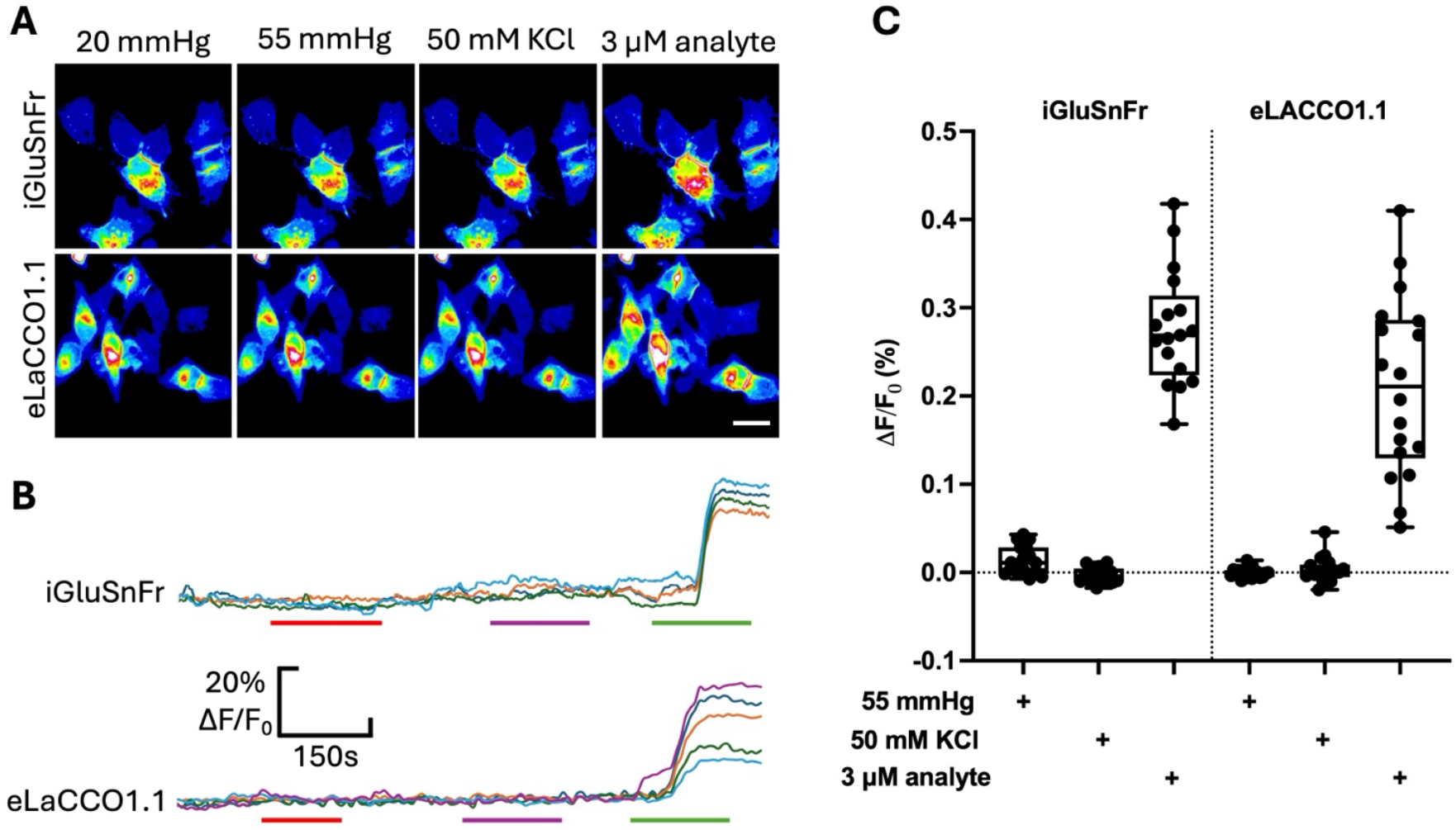
Parental HeLa DH cells do not release either glutamate or lactate in response to hypercapnia or depolarisation. A) Images of iGluSnFr or eLACCO1.1 fluorescence in HeLa cells. No change in fluorescence intensity was seen when the PCO_2_ of the aCSF was changed from 20 to 55 mmHg or when 50 mM KCl was applied. Scale bar 20 µm. B) Traces showing the plot of fluorescence changes (∧1F/F_0_) versus time for the images in A. Red bar, application of 55 mmHg PCO_2_; purple bar, 50 mM KCl; green bar, application of the relevant analyte at 3 µM; all stimuli applied from a baseline of 20 mmHg PCO_2_. C) Summary data showing that parental HeLa cells do not show fluorescence changes in response to CO_2_ or KCl. Data from 3 transfections each for iGluSnFr and eLACCO1.1.

**Figure 2.**
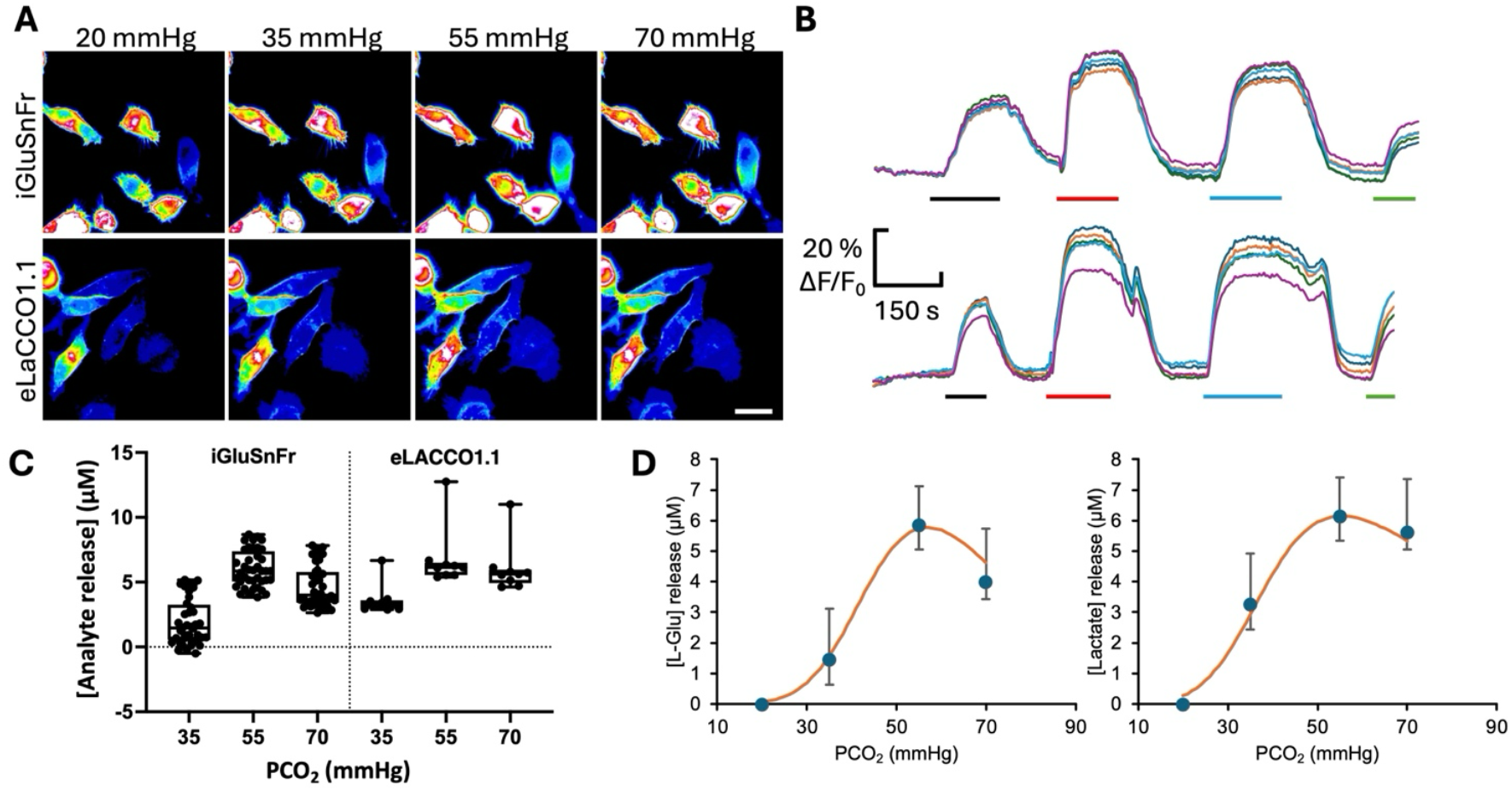
Connexin 50 is CO_2_ sensitive. **A)** Images of iGluSnFr or eLACCO1.1 fluorescence in HeLa cells that express Cx50 to show the progressive larger changes in fluorescence intensity when the PCO_2_ of the aCSF was changed from 20 to 35, 55 and 70 mmHg. Scale bar 20 µm. **B)** Traces showing the plot of fluorescence changes (ι1F/F_0_) for iGluSnFr (top) or eLACCO1.1 (bottom) versus time during application of 35 mmHg (black bar), 55 mmHg (red bar), 70 mmHg (light blue bar) and 3 µM analyte (green bar). **C)** Summary data plotted as box and whisker plots showing CO_2_ evoked glutamate and lactate release (from four independent transfections for each analyte). **D)** CO_2_-dependent release of glutamate and lactate from Cx50 expressing cells replotted as median with upper and lower quartiles. The continuous line is a modified Hill equation:

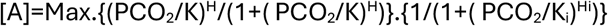

Where A is the analyte,, K and H are the atinity and Hill coeticient of the channel for opening by CO_2_, K_i_ and H_i_ are the atinity and Hill coeticient for inhibition of the channel by CO_2_, and Max (µM) is the asymptotically maximum released concentration of the analytes to CO_2_. The parameters for the curves are:

|  | Glu | Lac |
| --- | --- | --- |
| H | 6 | 5 |
| K (mmHg) | 46 | 42 |
| Max (μM) | 10 | 11.5 |
| K<sub>i</sub> (mmHg) | 70 | 70 |
| H<sub>i</sub> | 5 | 3 |

### The CO_2_-sensitivity of Cx50 depends on the carbamylation motif

We next tested whether the observed CO_2_ sensitivity of Cx50 hemichannels depended on K140 and K105 by individually mutating these residues to Gln. Gln cannot be carbamylated and, being neutral, will not be able to form a salt bridge with a carbamylated Lys residue. Application of 55 mmHg PCO_2_ solutions to HeLa cells expressing either Cx50^K140Q^ or Cx50^K105Q^ was ineffective at evoking glutamate release (Fig 3). As a positive control to demonstrate the presence of functional hemichannels, we used 50 mM KCl to depolarise the HeLa cells. This stimulus gave robust glutamate release showing that the mutated channels remained voltage sensitive and permeable to glutamate (Fig 3). We conclude that the action of CO_2_ in opening Cx50 hemichannels depends upon the carbamylation motif that we had previously identified in its structure.

**Figure 3.**
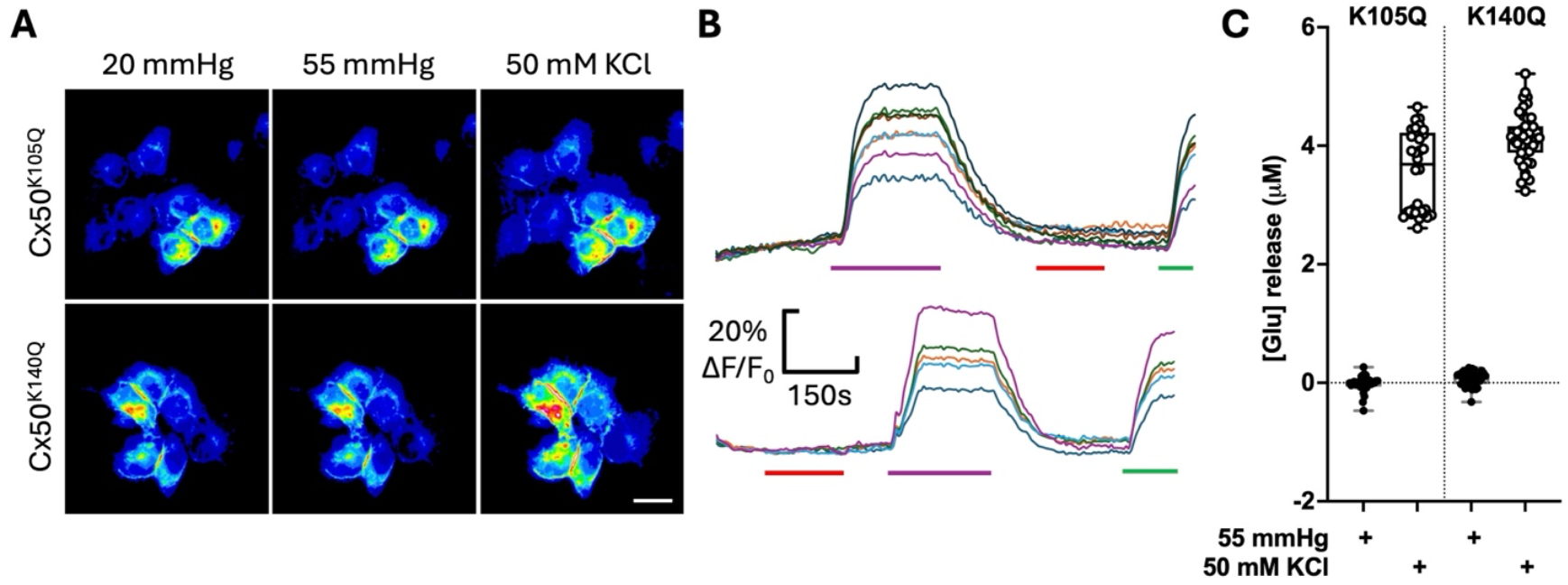
The CO_2_ sensitivity of Cx50 depends on the carbamylation motif residues K140 and K105. **A)** Images of iGluSnFr fluorescence showing that the mutations K105Q and K140Q abolish CO_2_-dependent glutamate release from HeLa cells expressing these mutant Cx50 hemichannels. A depolarising stimulus (50 mM KCl) was still able to evoke glutamate release demonstrating that the mutant hemichannels remain voltage dependent and permeable to glutamate. Scale bar 20 µm. **B)** Traces showing the plot of fluorescence changes (ι1F/F_0_) versus time for the images in A. Red bar, application of 55 mmHg PCO_2_; purple bar, 50 mM KCl; green bar 3 µM glutamate; all stimuli applied a baseline of 20 mmHg PCO_2_. **C)** Summary data from 4 independent transfections for K105Q and 5 independent transfections of K140Q showing that these mutations abolish the CO_2_ evoked glutamate release.

### Some cataract forming mutations of Cx50 alter its CO_2_-sensitivity

Cx50 plays a key role in the physiology of the lens and certain mutations cause cataract formation (Shiels *et al*., 1998; Berthoud & Ngezahayo, 2017; Berthoud *et al*., 2020; Shiels & Hejtmancik, 2021; Shi *et al*., 2022). We have examined whether three of these mutations V44A, V44E, W45S alter CO_2_ sensitivity (Fig 4). We found that both mutations of V44 abolished the CO_2_ sensitivity of Cx50 hemichannels, but the mutated channels remained voltage sensitive and permeable to glutamate. However, Cx50^W45S^ hemichannels remained CO_2_ sensitive but significantly less so than Cx50^WT^ (p<0.0001, Fig 4). Our results raise the possibility that loss or reduction of CO_2_ sensitivity of Cx50 could be a contributing cause to cataract formation.

**Figure 4.**
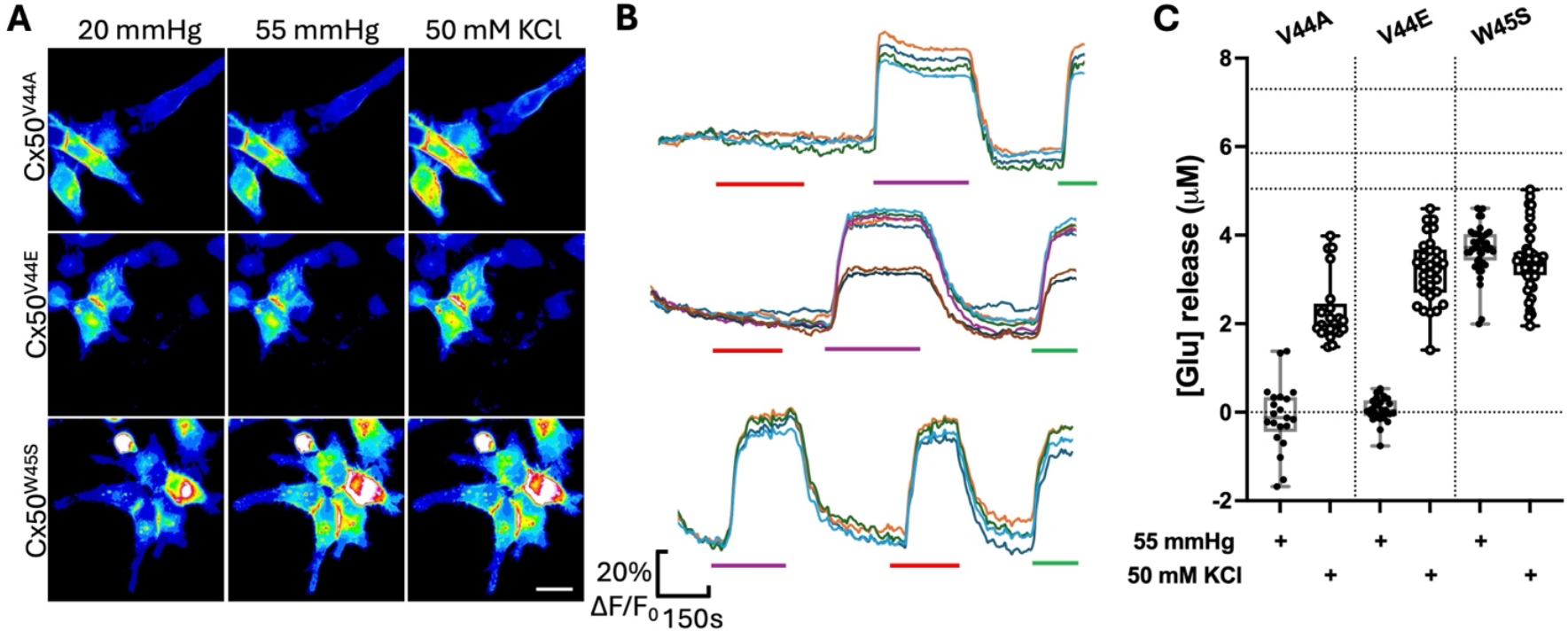
Some cataract forming mutations of Cx50 remove its CO_2_ sensitivity. **A)** Images of iGluSnFr fluorescence showing that the mutations V44E and V44A but not W45S abolish CO_2_-dependent glutamate release from HeLa cells expressing these mutant Cx50 hemichannels. A depolarising stimulus (50 mM KCl) is still able to evoke glutamate release demonstrating that the mutant hemichannels remain voltage dependent and permeable to glutamate. Scale bar 20 µm. **B)** Traces showing the plot of fluorescence changes (ΔF/F_0_) versus time corresponding to the images in A. Red bar, application of 55 mmHg PCO_2_; purple bar 50 mM KCl; green bar, 3 µM glutamate; all stimuli applied from a baseline of 20 mmHg PCO_2_. **C)** Summary data for: V44A, 4 independent transfections; V44E, 4 independent transfections; and W45S, 5 independent transfections. The dotted lines at top of y-axis show the values for the median CO_2_-evoked glutamate release with lower and upper quartiles for Cx50^WT^. Cx50^V44A^ and Cx50^V44E^ are insensitive to CO_2_, while Cx50^W45S^ releases significantly less glutamate to 55 mmHg than the wildtype channel (MW Test, p<0.0001).

To check whether the results of the mutations could be explained by altered membrane localisation, we stained the plasma membrane with DiO and examined colocalization of DiO with the mCherry tag on Cx50 for the WT channel and all mutated variants with confocal microscopy (Fig 5). This analysis showed that the Manders’ coefficients were the same for all Cx50 variants. Thus, the mutations did not alter membrane localisation of Cx50.

**Figure 5.**
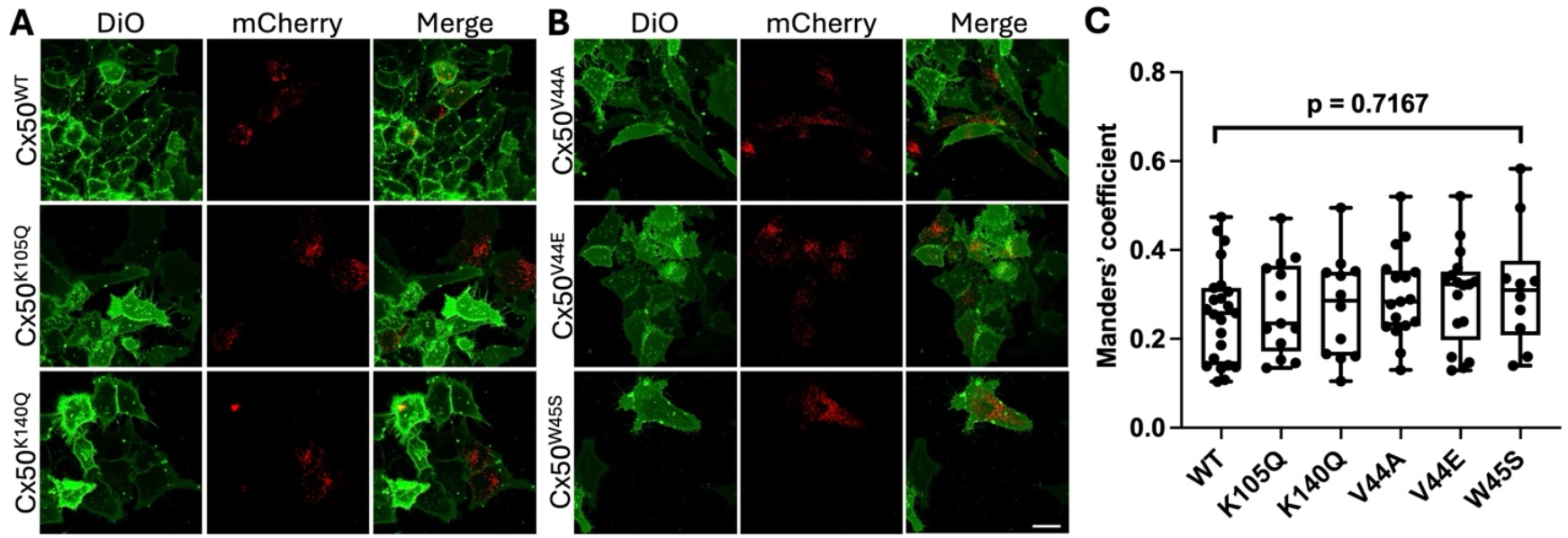
Carbamylation motif mutations and cataract forming mutations do not alter expression of Cx50 at the plasma membrane. **A,B)** Confocal images (single optical plane) showing membrane staining with DiO (green) and the mCherry C-terminal tag on the Cx50 variants (red). Scale bar 20 µm. **C)** Measurement of the Manders’ coeticient (the proportion of DiO fluorescence that colocalises with mCherry) for each Cx50 variant. These coeticients do not diter between the wild type and the mutants (Kruskal Wallis ANOVA, p=0.7167). Data from 3 independent transfections for each variant.

### The lens is CO_2_ sensitive

Cx50 is expressed in lens fibre cells and along with Cx46 is the major connexin subtype in this tissue. Given that Cx50 is CO_2_ sensitive, we might expect that this would endow the lens itself with CO_2_ sensitivity. We therefore used a dye loading assay to test this (Huckstepp *et al*., 2010b). We exposed acutely isolated whole lens to a membrane impermeant dye, FITC, at different levels of PCO_2_. Were Cx50 in the lens to open, then it should permit the entry of FITC into the fibre cells.

At 20 mmHg PCO_2_ (Cx50 hemichannels would be shut), we did not observe FITC loading in either differentiating or mature fibre cell types (Fig 6). FITC was frequently observed to accumulate between the plasma membranes, without entering cells. Trapping of Lucifer Yellow has also been observed between lens fibre cells (Suzuki-Kerr *et al*., 2022). By contrast at 55 mmHg PCO_2_ FITC fluorescence was observed in the cytosol of both differentiating and mature fibre cells (Fig 6).

**Figure 6.**
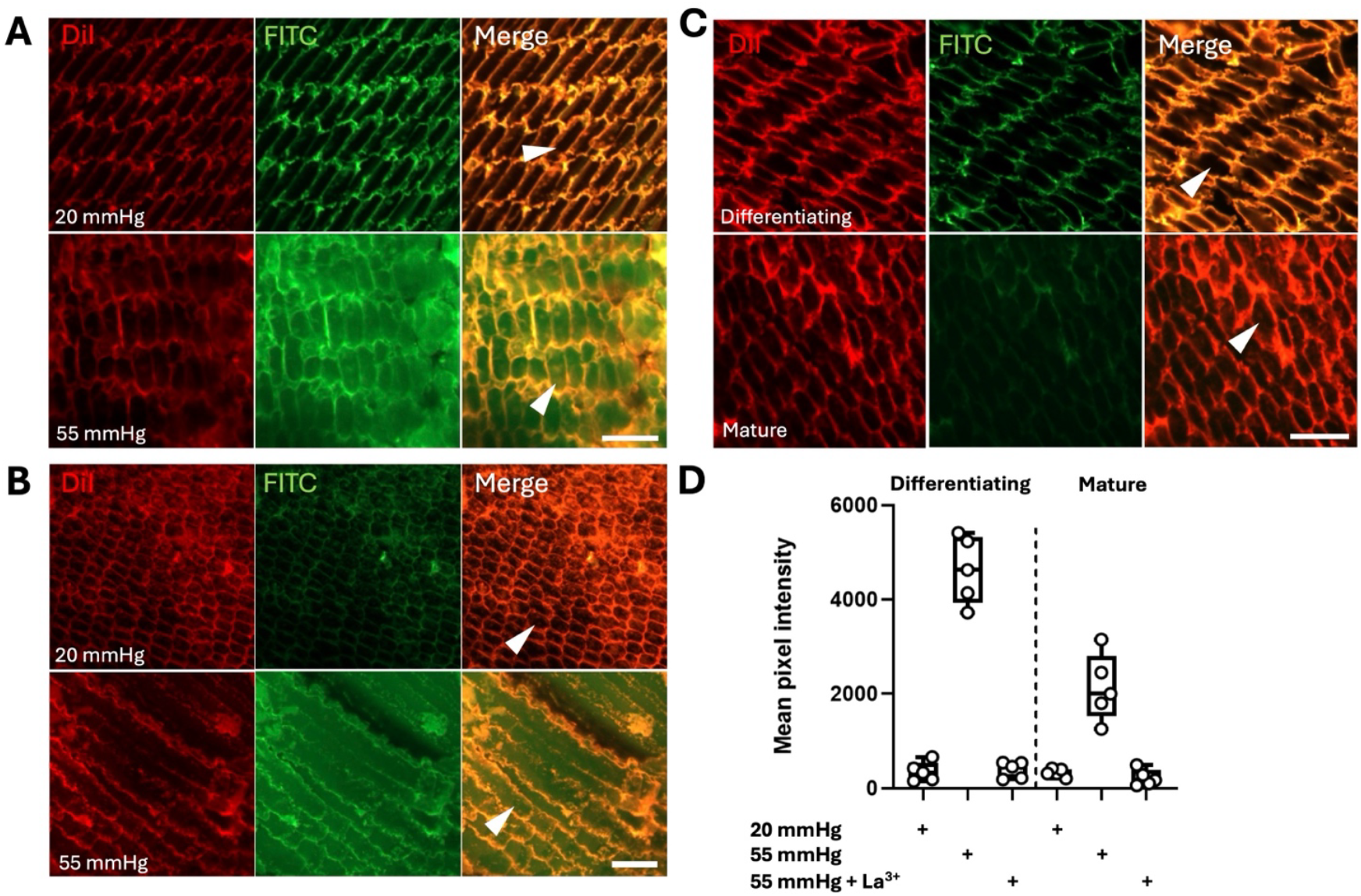
Hemichannel-mediated CO_2_-dependent dye loading into lens fibre cells. Diterentiating **(A)** and mature **(B)** fibre cells load with the membrane impermeant dye (FITC, green) in at a PCO_2_ of 55 mmHg but not at 20 mmHg. The sections of lens tissue have been stained with DiI (red) to label the membranes. Some FITC is present on the outside of the membrane in both conditions, but FITC is only present within the cytosol (arrow heads) at the higher level of PCO_2_. **C)** Dye loading at 55 mmHg PCO_2_ is completely blocked by the hemichannel inhibitor, La^3+^ (200 µM) -no FITCX is present in the cytosol of wither diterentiating or mature lens fibre cells (arrow heads). **D)** Summary graph showing for 5 lens preparations under each condition, the amount of FITC loading in fibre cells. Each individual data point represents the mean pixel intensity of at least 10 cells from a unique lens (n=5 lens). Kruskal Wallis ANOVA: Diterentiating fibres, p <0.0001, Mature fibres, p <0.0001. Post hoc Mann Whitney U tests: Diterentiating fibres; 20 vs 55 mmHg, p = 0.0056, 20 vs 55 mmHg + La^3+^, p = 0.7682, Mature fibres; 20 vs 55 mmHg, p = 0.0005, 20 vs 55 mmHg + La^3+^, p = 0.9492. Scale bars 25 µm (A), 40 µm (B), 25 µm (C).

To test whether the FITC entry might occur via a hemichannel, we used the blocker La^3+^ which blocks many types of hemichannel (connexin and pannexin) but does not block voltage-gated Na^+^, K^+^ or Ca^2+^ channels. La^3+^ completely blocked FITC entry during 55 mmHg (Fig 6), indicating that this occurred via a hemichannel. As Cx50 hemichannels are the only CO_2_ sensitive large-pored channel that could plausibly permit the entry of FITC, it is very likely that these hemichannels are the conduit for FITC entry.

### Lens fibre cells exhibit CO_2_ dependent Ca^2+^ entry

Given that lens microcirculation depends on ion influxes in lens fibre cells, we sought to gain evidence for CO_2_-dependent ion entry into lens fibre cells. As it is much easier to measure intracellular Ca^2+^ than Na^+^ and connexin hemichannels are permeable to both Na^+^ and Ca^2+^ ions, we loaded acute lens slices with Fluo-4 to test whether changes in PCO_2_ might trigger a Ca^2+^ influx. Using a PCO_2_ of 20 mmHg as a baseline, we demonstrated that increases to either 35 mmHg or 55 mmHg triggered a very substantial influx of Ca^2+^ that could be blocked by pre-treatment with La^3+^ (Fig 7). This indicates that opening of a CO_2_ sensitive hemichannel, most likely Cx50, provides a Ca^2+^ influx. Crucially, the CO_2_ sensitivity of this influx matches the dose response of Cx50 that we established in Fig 2 and shows that even at the likely resting PCO_2_ of aqueous humour, Cx50 hemichannels in lens fibre cells will be partially open.

**Figure 7.**
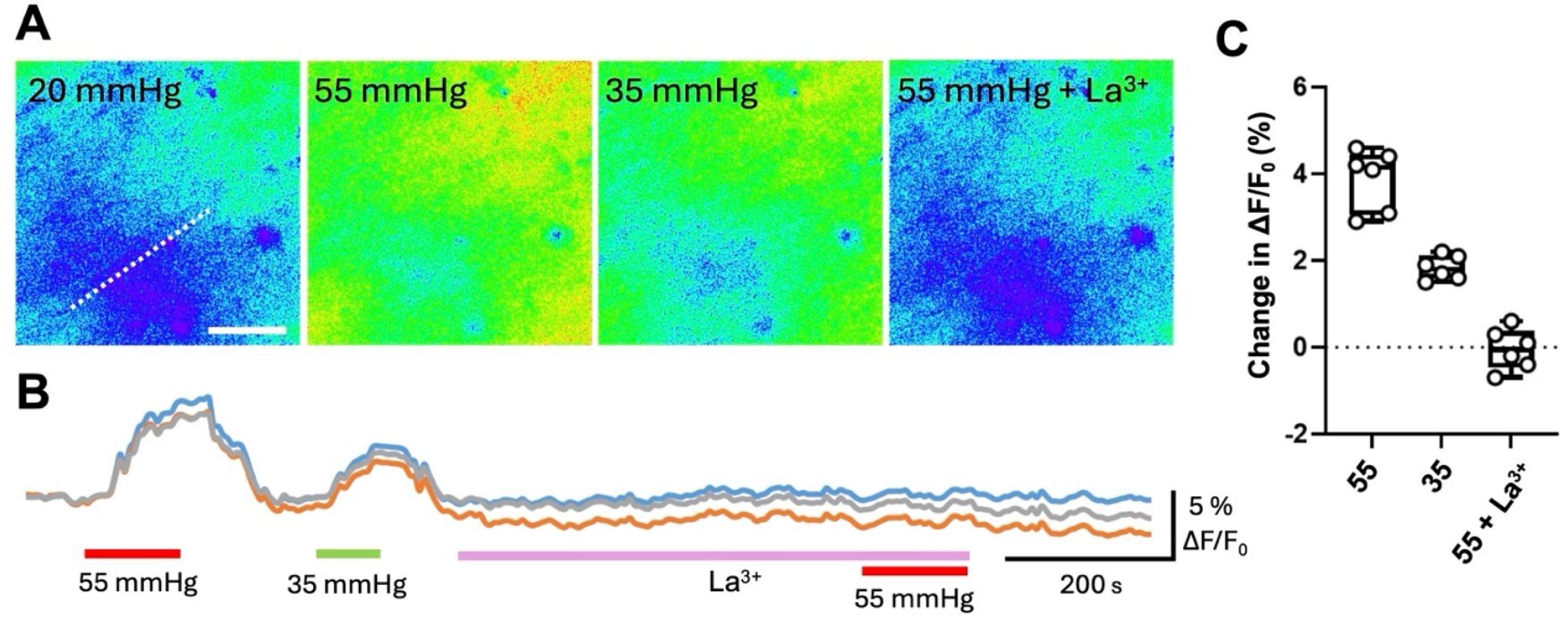
Hemichannel mediated CO_2_ dependent Ca^2+^ influx into lens fibre cells. **A)** Sample images of Fluo-4 loaded lens slices. The dotted line is parallel to the long axis of the fibre cells in the slice. A large increase of fluorescence was seen at 55 and 35 mmHg PCO_2_ compared to the baseline (20 mmHg). The increase in fluorescence evoked by 55 mmHg PCO_2_ was blocked by 200 µM La^3+^. Scale bar, 50 µm. **B)** Quantification of the fluorescence changes evoked by 55 and 35 mmHg and the blocking effect of La^3+^ on the increase in fluorescence. **C)** Summary data from 6 different lens slices showing the changed in normalised fluorescence under the three conditions. Friedman 2-way ANOVA: p = 0.0278. Wilcoxon matched-pairs signed rank test: 55 vs 35 mmHg PCO_2_, p = 0.0313.

### Lens fibre cells respond to glutamate via NMDA receptors

A curious feature of Cx50 hemichannels is that they are impermeable to ATP, but permeable to both glutamate and lactate (Lovatt *et al*., 2025). This led us to question whether a glutamate efflux via the opening of Cx50 hemichannels might be physiologically important in lens. There are several reports of AMPA and NMDA receptors being expressed on lens fibre cells (Farooq *et al*., 2012; Bhattacharyya *et al*., 2014; Frederikse *et al*., 2016), but the source of any activating glutamate and their roles in the physiology of lens remain unknown.

We therefore tested whether glutamate might evoke Ca^2+^ signals in lens fibre cells. Using Fluo-4 loaded acute lens slices, we found that glutamate evoked robust increases in intracellular Ca^2+^. These could be blocked by the general glutamate-receptor antagonist kynurenic acid and the NMDA receptor selective antagonist D-AP5, but not by the AMPA receptor selective antagonist, CNQX (Fig 8). The glutamate responses were thus mediated via NMDA receptors. Note that we cannot exclude a role for the AMPA receptor in mediating ion influx, but as the AMPA receptor in lens undergoes post-transcriptional Q/R editing (Farooq *et al*., 2012) it is impermeable to Ca^2+^ and we would therefore not be able to observe any contribution from this receptor via Fluo-4 imaging. The glutamate responses were not blocked by La^3+^, indicating that they were independent of any hemichannel activity (Fig 8).

**Figure 8.**
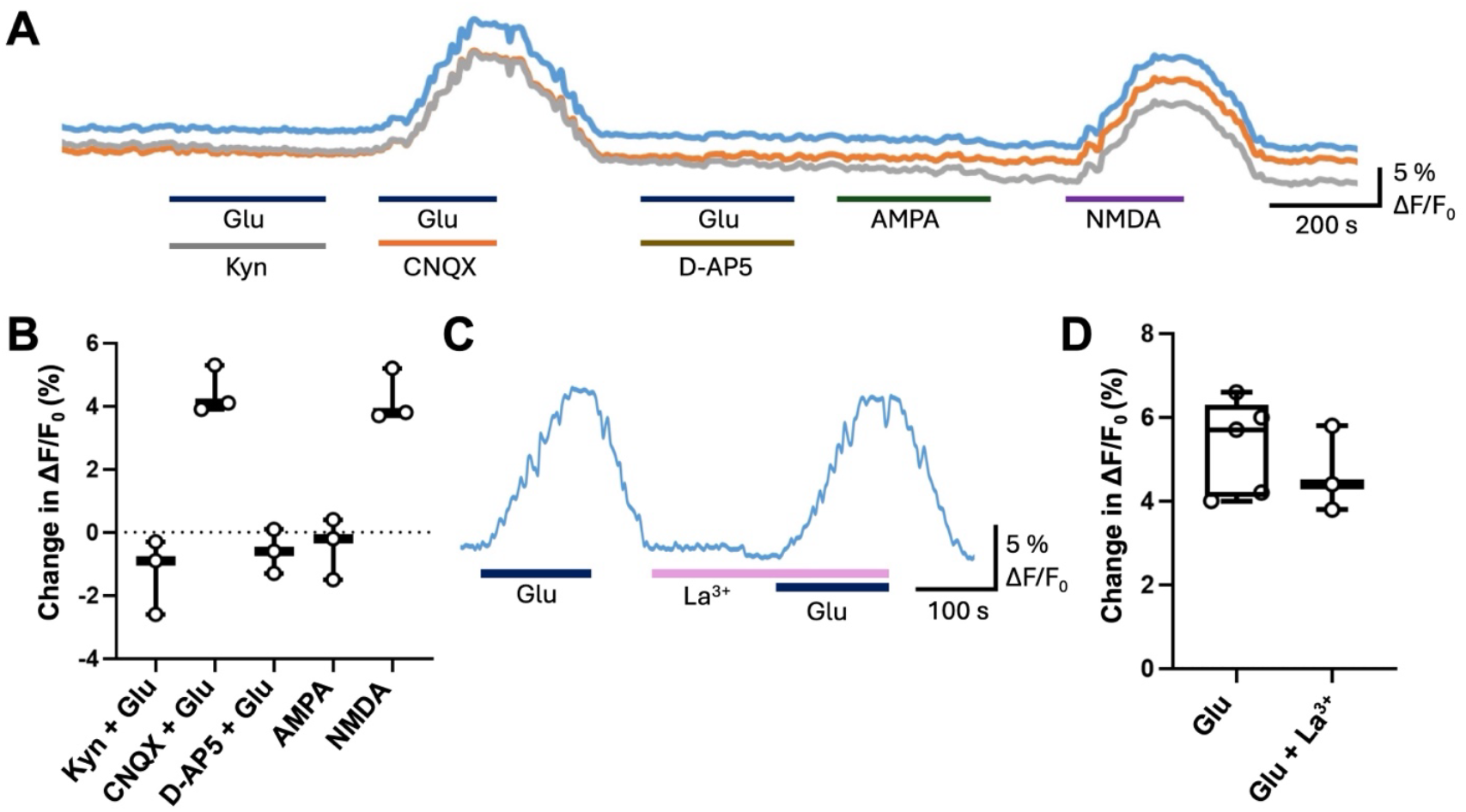
Glutamate evoked Ca^2+^ signals in lens fibre cells are mediated by NMDA receptors. **A)** Traces of normalized Fluo-4 fluorescent showing the response to 200 µM L-glutamate in presence of 1 mM kynurenic acid, 10 µM CNQX or 100 µM D-AP5, and the selective ionotropic glutamate receptor agonists AMPA (5 µM) and NMDA (60 µM). B) Summary data for the responses to glutamate receptor agonists for n=3 lens slices. Friedman 2-way ANOVA: p = 0.0053. **C, D)** The responses to glutamate are not dependent on hemichannel activation and are unatected by 200 µM La^3+^, n=5 lens slices for Glu and n=3 lens slices for Glu + La^3+^.

### CO_2_-evoked glutamate release contributes to the CO_2_ dependent Ca^2+^ influx in lens fibre cells

Given that Cx50 hemichannels permit the CO_2_-dependent release of glutamate, we asked whether the Ca^2+^ influx triggered by 55 mmHg PCO_2_ might be at least partially mediated via NMDA receptors. We therefore loaded lens fibre cells with Fluo-4 and measured first the change intracellular Ca^2+^ evoked by the PCO_2_ challenge and then applied D-AP5 prior to the PCO_2_ challenge to see whether NMDA receptor antagonism might reduce the Ca^2+^ signal. We found that D-AP5 reversibly reduced the Ca^2+^ influx by roughly half (Fig 9). This implies that during elevated PCO_2_ the opening of Cx50 hemichannels permits not just the influx of Ca^2+^ but also the efflux of glutamate which can then activate NMDA receptors to provide an additional Ca^2+^ influx. The lens fibre cells are thus a source of the glutamate that activates the receptors on their cell surface.

## Discussion

Cx50 is the fifth connexin, after Cx26, Cx30, Cx32 and Cx43, that is now known to be CO_2_ sensitive. Significantly, it is the second alpha connexin alongside Cx43 documented to have this property. All 5 connexins share a carbamylation motif that was originally identified in Cx26 and closely related beta connexins. The mechanism of CO_2_ dependent hemichannel opening in Cx43 involves 4 Lys residues: K105, K109, K144 and K234 (Dospinsecu *et al*., 2025). Cx50 lacks K109 (this residue is an Arg) and it is interesting that mutation of either K105 or K140 in Cx50 is sufficient to abrogate CO_2_ sensitivity. In Cx43 at least 2 Lys residues must be mutated to abolish CO_2_ sensitivity (Dospinsecu *et al*., 2025). Thus, the mechanism of CO_2_ sensitivity in Cx50 is somewhat simpler than in the related Cx43 and more like that proposed for Cx26 (Meigh *et al*., 2013; Brotherton *et al*., 2022; Brotherton *et al*., 2024).

**Figure 9.**
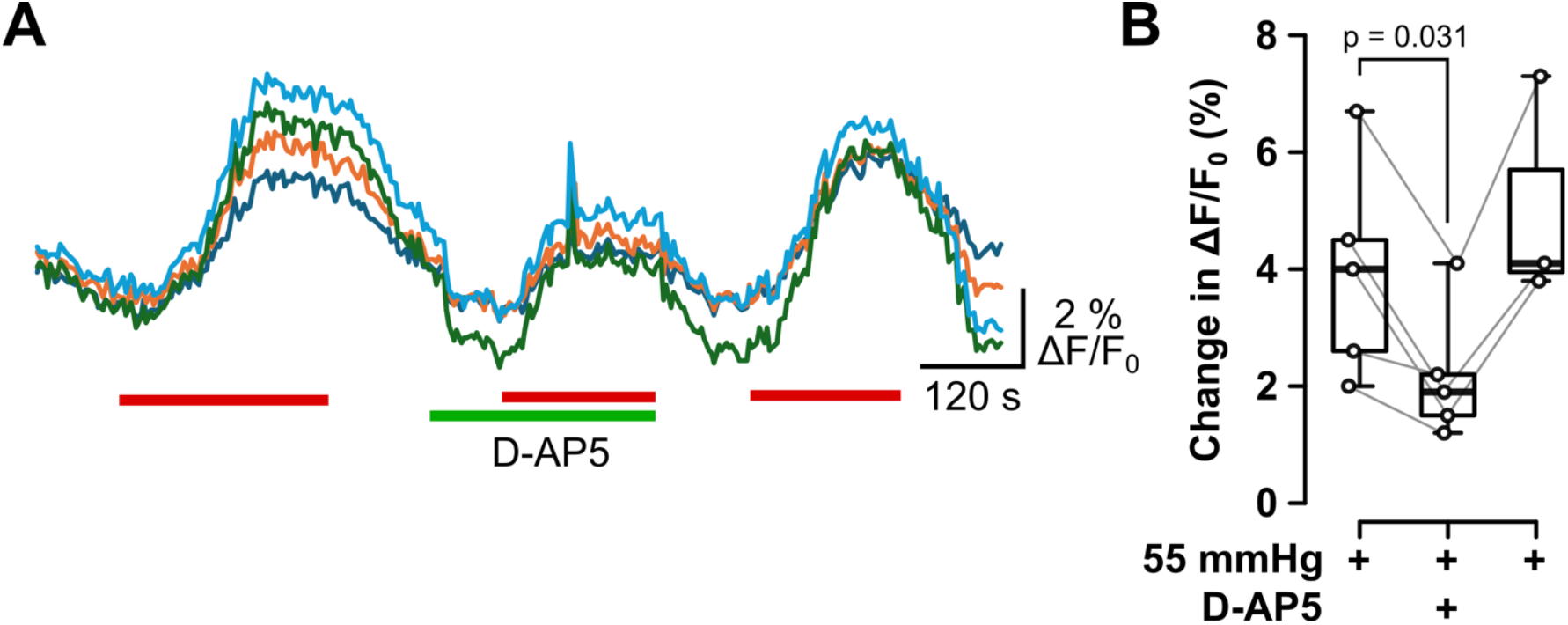
Activation of NMDA receptors during a CO_2_ challenge partially contributes to the elevation of intracellular Ca^2+^ in lens fibre cells. **A)** Traces from ROIs on a Fluo-4 loaded lens slice showing responses to 55 mmHg PCO_2_ (red bars) before, during and after application of 100 µM D-AP5 (green bar). **B)** Summary graph showing etect of 100 µM D-AP5 on change in intracellular Ca^2+^ evoked by 55 mmHg PCO_2_; n=5 lens for 55 mmHg and 55 mmHg + D-AP5, n=3 lens for wash after 55 mmHg + D-AP5. Wilcoxon matched pairs signed rank test, p=0.031, 55 mmHg vs 55 mmHg + D-AP5.

### Could Cx50 hemichannels act as homeostatic regulators of lens microcirculation?

The properties of Cx50 hemichannels appear to be matched to the physiology of the lens. Firstly, Cx50 hemichannels are significantly open at a PCO_2_ of 35 mmHg. PCO_2_ of aqueous humour is generally observed as being a little lower than that measured in plasma (Salit, 1930; Krupin *et al*., 1980; Sharma *et al*., 1983) and is in the range 35-40 mmHg. According to our data therefore, Cx50 hemichannels would be expected to be partially open and provide a continual source of Na^+^ ions to power the microcirculation. Secondly the degree of opening of Cx50 will be proportional to PCO_2_ over the relevant physiological range. We propose the hypothesis that CO_2_, acting via Cx50, could mediate homeostatic control of microcirculation. Under this hypothesis, were PCO_2_ to rise above its normal resting level, it would imply insufficient microcirculation and waste removal. As a consequence of the increased PCO_2_, Cx50 hemichannels would open more and allow a greater Na^+^ influx leading to faster microcirculation and a corrective adaptive response. Conversely, if PCO_2_ were to fall below the normal resting level, it would imply that the microcirculation was too fast. Cx50 hemichannels would close in response to the lowered PCO_2_, reducing the Na^+^ influx and hence the rate of microcirculation once again leading to a corrective adaptive response.

Thirdly, Cx50 hemichannels are permeable to lactate. To maintain transparency of the lens, as lens fibre cells mature, they lose their organelles including mitochondria (Bassnett, 2002). This means that they become dependent on ATP supply from more metabolically active cells via gap junctions and from glycolysis. Lactate is the end product of glycolysis and thus the permeability of Cx50 to lactate potentially provides an effective way of allowing efflux from the fibre cells of this end product.

### Supporting evidence for the proposed homeostatic role of Cx50 hemichannels

We have shown that the lens is indeed CO_2_ sensitive with characteristics consistent with the properties of Cx50 hemichannels. Lens fibre cells load with FITC in a CO_2_ sensitive manner, and we recorded a CO_2_-dependent Ca^2+^ influx that is likely through Cx50, as it matched the CO_2_ dose dependence of the Cx50 hemichannel and, like the FITC-loading, could be blocked by La^3+^. While this does not definitively demonstrate that lens CO_2_ sensitivity depends on Cx50, the balance of probabilities makes this likely.

Our evidence gives some support to Cx50 hemichannels acting as a homeostatic regulator of lens microcirculation. Firstly, there was an influx of Ca^2+^ into fibre cells at a PCO_2_ of 35 mmHg, this supports our speculation that Cx50 hemichannels in lens fibre cells may be partially open at the resting PCO_2_ typical of aqueous humour. Secondly, the Ca^2+^ influx increases with increasing PCO_2_ supporting the second component of our hypothesis. Thirdly, our evidence ties together a curious property of the Cx50 hemichannel, its permeability to glutamate, with the existence of ionotropic glutamate receptors on lens fibre cells. We found that the NMDA receptor antagonist, D-AP5, partially reduced the CO_2_-induced Ca^2+^ influx into the fibre cells. This is most simply explained by the opening of Cx50 hemichannels not only allowing the influx of Ca^2+^ but the efflux of glutamate which can then activate NMDA receptors to give an additional component of Ca^2+^ influx. We note that both Cx50 hemichannels and NMDA receptors are highly permeable to Na^+^, so will act as a source (presumably via gap junction coupling) of Na^+^ for the Na^+^/K^+^ ATPases in the lens epithelial cells. Furthermore, glutamate released via Cx50 hemichannels will most probably also activate the AMPA receptors present on the fibre cells, even though we cannot directly observe this via Ca^2+^ imaging.

While many investigators have queried the possible identity of the channel that provides the Na^+^ inward leak current to power microcirculation, our data suggests that this Na^+^ influx may originate from multiple channels: Cx50 as the trigger, with downstream activation via glutamate release of the NMDA receptor, and very likely the AMPA receptor, providing additional sources of Na^+^ influx. There is also evidence that Cx46 hemichannels could contribute to a Na^+^ leak current in lens fibre cells (Ebihara *et al*., 2014). It is important to note that we have not directly measured lens microcirculation and that definitive testing our hypothesis requires demonstration that the rate of microcirculation in the lens can be altered in a CO_2_-dependent manner that is consistent with the properties of Cx50 hemichannels and the involvement of glutamate receptors.

### Pathological mutations of Cx50 alter its CO_2_ sensitivity

If Cx50 hemichannels really are a key regulator of the microcirculation in the lens, at least some cataract forming mutations of Cx50 should alter its CO_2_ sensitivity. We have previously shown that a subset of pathological mutations of Cx26, Cx32 and Cx43 that respectively cause keratitis ichthyosis deafness syndrome (KIDS), X-linked Charcot Marie Tooth Disease (CMTX) and oculodentodigital dysplasia (ODDD) abolish the CO_2_ sensitivity of these hemichannels (Meigh *et al*., 2014; de Wolf *et al*., 2016; Cook *et al*., 2019; Butler & Dale, 2023; Dospinsecu *et al*., 2025). Our observation that some cataract-forming mutations of Cx50 also modify its CO_2_ sensitivity therefore fits this pattern. Our observation is consistent with, but does not substantiate, the hypothesis that the loss of CO_2_ sensitivity of Cx50 contributes to cataract formation. As Cx50^W45S^ retains reduced CO_2_-sensitivity yet also causes cataracts, loss of other properties of Cx50 must also contribute to cataract formation. One might imagine that any mutations that alter the permeability of Cx50 hemichannels to glutamate might also cause cataracts if our hypothesis is correct. Clearly, examination of further cataract mutations will be required to understand any potential role that loss of CO_2_ sensitivity might have in the aetiology of cataract formation. In addition, testing whether mutations that directly target CO_2_ sensitivity (e.g. K140Q) are sufficient to trigger cataract formation would be a more rigorous test of this hypothesis.

